# miR-125b-5p impacts extracellular vesicle biogenesis, trafficking, and EV subpopulation release in the porcine trophoblast by regulating ESCRT-dependent pathway

**DOI:** 10.1101/2023.04.10.536278

**Authors:** Maria M. Guzewska, Krzysztof J. Witek, Elżbieta Karnas, Michał Rawski, Ewa Zuba-Surma, Monika M. Kaczmarek

## Abstract

Intercellular communication is a critical process that ensures cooperation between distinct cell types at the embryo–maternal interface. Extracellular vesicles (EVs) are considered to be potent mediators of this communication by transferring biological information in their cargo (e.g. miRNAs) to the recipient cells. miRNAs are small non-coding RNAs that affect the function and fate of neighboring and distant cells by regulating gene expression. Focusing on the maternal side of the dialog, we recently revealed the impact of embryonic signals, including miRNAs, on EV-mediated cell-to-cell communication. In this study, we show the regulatory mechanism of the miR-125b-5p ESCRT-mediated EV biogenesis pathway and the further secretion of EVs by trophoblasts at the time when the crucial steps of implantation are taking place.

To test the ability of miR-125b-5p to influence the expression of genes involved in the generation and release of EV subpopulations in porcine conceptuses, we used an ex vivo approach. Next, in silico and in vitro analyses were performed to confirm miRNA–mRNA interactions. Finally, EV trafficking and release were assessed using several imaging and particle analysis tools.

Our results indicated that conceptus development and implantation are accompanied by changes in the abundance of EV biogenesis and trafficking machinery. ESCRT-dependent EV biogenesis and the further secretion of EVs were modulated by miR-125b-5p, specifically impacting the ESCRT-II complex (via VPS36) and EV trafficking in primary porcine trophoblast cells. The identified miRNA–ESCRT interplay led to the generation and secretion of specific subpopulations of EVs.

miRNA present at the embryo–maternal interface governs EV-mediated communication between the mother and the developing conceptus, leading to the generation, trafficking, and release of characteristic subpopulations of EVs.

## 2. INTRODUCTION

Cell communication between a developing embryo and the maternal endometrium is based on a sensitively timed exchange of a wide range of biologically active molecules, which is crucial for the molecular processes that allow for a successful pregnancy [1]. Extracellular vesicles (EVs) are recognized as crucial mediators of cell-to-cell signal transmission. The EV-mediated exchange of signals at the embryo–maternal interface is now considered to be a fundamental element of successful reproduction, promoting embryo implantation and assisting in pregnancy [2]. Biological pathways leading to the formation and secretion of EV subtypes are under cellular control and can be governed by different internal and external factors [3]. Based on the latest guidelines, heterogeneous populations of EVs can be divided into the following three main groups based on their physical size range: small EVs, medium/large EVs, and apoptotic bodies [4]. Small EVs, formed as intraluminal vesicles (ILVs), are created during the maturation of early endosomes to late endosomes. By the inward budding of the membrane, late endosomes change into multivesicular bodies (MVBs) filled with ILVs. The endosomal sorting complex required for transport (ESCRT) machinery plays an important role in the formation of MVBs and ILVs [5]. The ESCRT pathway is not only a key regulator of MVB biogenesis, but it is also in charge of ubiquitinylated protein sorting into EVs, which is executed by four ESCRT complexes. The ESCRT is comprised of an ancient system, conserved from yeast to mammals, for the membrane remodeling and scission of approximately 20 proteins that assemble into ESCRT-0, -I, -II, and -III complexes. These complexes are supported by associated proteins like the programmed cell death 6 interacting protein (ALIX) and ATPase – vacuolar protein sorting 4 (VPS4) [6]. Importantly, vesicles can also be generated through ESCRT-independent pathways. During biogenesis, vesicles gain various surface markers found at the double membrane, such as tetraspanins (e.g., CD63, CD81, CD9), also used to target recipient cells and uptake [7,8,9,10]. Tetraspanins, such as CD63, have also been shown to be important in ESCRT-independent loading into small EVs, while their interaction with other transmembrane/cytosolic proteins and lipids organizes the membrane into tetraspanin-enriched domains [11]. The model of biogenesis is critically related to the physiologic function of EVs and their selected cargo, as well as the pathway of secretion governed by small Rab GTPases [12]. Rab GTPases are widely considered to be cellular organelle markers, since each Rab protein regulates a distinct transport step related to endosome generation, MVB maturation, ILV formation, and EV secretion [12]. Because of the protection by the double membrane, EVs can transport a wealth of cargos, including not only proteins but also lipids and nucleic acids – microRNAs (miRNAs) and mRNAs. Importantly, the autocrine, paracrine, or endocrine functions of the delivered EVs depend on the content of the packed cargo [13].

Interestingly, EVs can transfer miRNAs to acceptor cells to downregulate the expression of target genes. miRNAs are endogenous, small RNAs with an average of 22 nucleotides in length, which can inhibit mRNA translation and/or induce mRNA transcript degradation by binding mainly to the 3’ untranslated region (3’UTR) [14]. MiRNA are involved in a variety of biological processes, including cell proliferation, cell differentiation, and apoptosis [14]. Importantly, a single miRNA can target hundreds of different target genes and a single gene can be regulated by several different miRNAs [15,16]. Based on current knowledge, a large number of miRNAs have been identified as important regulatory molecules during early pregnancy in mammals, including pigs [17,18,19,20,21,22], sheep [23,24], and cattle [25]. Our previous research showed that embryonic [18] as well as endometrial [17] miRNAs are important players during early pregnancy in pigs, having the potential to affect gene expression at the embryo–maternal interface, as well as trophoblast cell functions [20]. A plethora of miRNAs were also found to be carried by EVs isolated form uterine flushings during early pregnancy [19]. Our recent studies have also indicated possible miRNA-mediated regulation of EV biogenesis and trafficking at the maternal site during pregnancy [26]. Nevertheless, many aspects of the way in which miRNAs encapsulated into EVs can regulate EV-mediated cell-to-cell communication is still not completely understood.

In this work, we sought to analyze the impacts of miR-125b-5p, encapsulated in EVs [18,19] and upregulated in porcine conceptuses after day 16 of pregnancy [18], on EV biogenesis pathways and their later trafficking and subpopulations secretion. To test the ability of miRNA to influence the expression of genes involved in the generation and release of EV subpopulations in porcine conceptuses, we used ex vivo and in vitro approaches. We identified inverse expression patterns of miR-125b-5p and genes coding ESCRT complex molecules and their associated proteins. In silico analysis confirmed miR-125b-5p targets among ESCRT complex genes, which was further proved by the decreased abundance of mRNA after the delivery of miR-125b-5p to porcine trophoblast cells (pTr) in vitro. Furthermore, miR-125b-5p occurred as a negative regulator of VPS36 recruitment at pTr cell borders, leading to the increased cellular abundance of CD63 tetraspanin and release of CD63^+^ EVs.

## 3. MATERIALS AND METHODS

### 3.1. Animals

Crossbred (Hampshire × Duroc) of similar age (7–8 months) and weight (140 ± 150 kg), after their second natural estrus, were artificially inseminated and again 24 h after onset of estrus (first day of pregnancy [DP]). Samples were collected post-mortem at local abattoir. Uterine horns isolated from pregnant animals were flushed with 40 mL 0.01 M PBS (Phosphate-buffered saline; pH 7.4) in order to recover conceptuses. The day of pregnancy was confirmed by size and morphology of conceptuses, as previously described [18]. For mRNA isolation embryos were collected on days 11– 12 (n = 8), 15–16 (n = 8), and 17–19 (n = 6), immediately frozen in liquid nitrogen and stored at −80°C until further use. Conceptuses from day 16 were placed into Dulbecco’s Modified Eagle Medium/Nutrient Mixture F-12 Ham medium (DMEM/F12; Sigma - Aldrich, USA) supplemented with 1% (v/v) penicillin/streptomycin (P/S; Sigma - Aldrich) and transported to the laboratory in order to isolate porcine primary trophoblast (pTr) cells for the following in vitro experiments.

### 3.2 Isolation and culture of porcine primary trophoblast cells

Porcine primary trophoblast cells were isolated according to the method established previously [27]. Conceptuses were washed three times in DMEM/F12 medium and then digested with 0.25% trypsin (Biomed, Lublin, Poland) for 30 min in 38°C. The suspension was filtered through sieve mesh filled with fresh medium containing 10% Newborn Calf Serum (NCS, Sigma - Aldrich, USA) in order to inactivate trypsin and then centrifuged at 200 × g for 10 min at 8°C. After final centrifugation, the obtained pellet was suspended in culture medium (DMEM/F12 containing 10% NCS and P/S). For mRNA/miRNA abundance evaluation, pTr cells were seeded on collagen-I coated 6-well plate (0.5 × 10^6^ / per well; Corning, USA); for EVs isolation on cell culture flask T-75 (1.5 − 2 × 10^6^/ per flask; Eppendorf, Hamburg, Germany) and for immunostaining on Cell Imaging Cover Glasses (0.1 × 10^6^ / per chamber; Eppendorf, Hamburg, Germany).

### 3.3. miRNA transfection

pTr cells were transfected with miRNAs after reaching 80% confluence. MiR-125b-5p (MIMAT0000423) or control mimic (Life Technologies, USA, Supplementary Table 1) were delivered to the cell culture using Lipofectamine RNAiMAX (Life Technologies, USA). Cells were washed twice with PBS and then incubated in DMEM-F12 with mimic-lipofectamine complexes for 12 h. The dose of 50 nM was selected based on a dose-dependent experiments [18,20]. After transfection medium was changed to culture medium and cells were cultured for additional 36 h at 37°C in a humidified atmosphere of 95 % air to 5 % CO_2_. Afterwards, culture medium was removed and trophoblast cells were washed in PBS, lysed in TRI Reagent (Invitrogen, USA) and subjected to total RNA isolation.

### 3.4. Immunofluorescent staining and confocal imaging

To investigate the effect of miRNAs on the cellular distribution of proteins belonging to the EV biogenesis pathway, immunofluorescent staining analysis was performed. pTr cells were cultured in Cell Imaging Coverglasses (Eppendorf, Germany) and subjected to miRNA transfection as described above. Then, pTr cells were treated as follows: 1) fixed immediately (n = 4); or 2) subjected to baculovirus treatment (CellLight™ Late Endosomes-GFP, BacMam 2.0, Invitrogen; dilution 1:50; incubation for 36 h at 37°C in a humidified atmosphere of 95 % air to 5 % CO_2_; n = 4). After fixation in 4% buffered formalin (Neutral Phosphate Buffered Formalin Fixatives, Leica, Germany), cells were permeabilized in 0.25% Triton X-100 and blocked in Sea Block Blocking Buffer (1 h; Thermo Fisher Scientific, USA). Overnight incubation with primary antibodies (anti-VPS36, anti-CD63, and anti-Rab7B) was followed by the incubation (1 h) with secondary antibodies conjugated with Cy3 or Alexa Fluor 488 dye (Supplementary Table 2). At the end of incubation, cells were treated with DAPI solution (0.1 mg/mL; Sigma - Aldrich, USA) according to the manufacturer protocol. Negative control staining was performed without primary antibodies and F-actin filaments were visualized using phalloidin (200 units/mL; Thermo Fisher Scientific, USA, Supplementary Table 2) conjugated with Alexa Fluor 488 (Supplementary Fig. 1A) or Alexa Fluor 594 (Supplementary Fig. 1B). Images were captured using confocal Laser Scanning Microscope (LSM800; Carl-Zeiss, Germany) equipped with Airyscan super-resolution module using 63x/1.4 NA c-apochromat objective. Colocalization analysis was performed as described previously [26]. Briefly, ten images of trophoblast cell colonies, showing at least 10 well-visible cells were captured. Regions of interests (ROIs) were designated by the eye-based evaluation of the quantity and quality of the pTr cells in monolayer. Localization of both protein markers (VPS36 and CD63) were determined at the edges of cells (described as cell-cell borders) and in the whole cell (described as whole cell; measurements included cell-cell borders). Next, VPS36 and CD63 fluorescence intensity as well as their colocalization coefficient (CC) were assessed [28]. CC indicates the relative number of colocalized pixels in relation to the total number of pixels above the constant image background-noise threshold value. Image analysis was done using ZEN Blue Pro 2.6 software (Carl-Zeiss, Germany).

### 3.5. EV isolation after miR-125b-5p transfection

pTr cells were seeded onto T-75 culture flasks (n = 5) and cultured until reaching 80% confluence. Transfection with miRNA was performed as described above. After 12 h of incubation with miRNA and lipofectamine complexes, cells were washed twice with PBS without Ca^2+^ and Mg^2+^ (Lonza, USA) and medium was changed to EV-depleted medium (DMEM F12 + 10% NCS, centrifuged at 100 000 × g, overnight at 4°C). After additional 36 h of culture, conditioned medium was collected in order to isolate secreted pTr - EVs. Collected medium was subjected to stepwise centrifugation at 4°C (156 × g, 10 min; 2 000 × g, 10 min; 10 000 × g, 30 min) to eliminate dead cells and cell debris. Final supernatant was passed through 0.22 µm filters and 40 mL of filtrate (from each: miR-125b-5p and control mimic) was ultracentrifuged twice in sterile 10.4 mL polycarbonate bottles at 100 000 × g for 70 min at 4°C (Optima L-100 XP Ultracentrifuge, 90 Ti Fixed-Angle Titanium Rotor; Beckman Coulter). The final EV pellet was suspended in 50 µl of PBS without Ca^2+^ and Mg^2+^ (Lonza, USA) and stored at −80°C for further analysis.

### 3.6 Cryo-electron microscopy of secreted pTr - EVs

To characterize secreted pTr - EVs we used cryo-electron microscopy (cryo-EM), a powerful tool for assessing morphology of EVs, which preserves membranes in a close to native state. Sample solution (~ 3 μl) was applied on freshly glow discharge TEM grids (Quantifoil R2/1, Cu, mesh 200) and plunge-frozen in liquid ethane using Vitrobot Mark IV (Thermo Fisher Scientific). The following parameters were set: humidity - 100%, temperature - 4°C, blot time - 2 sec. Frozen grids were kept in liquid nitrogen until clipping and loading into the microscope. Cryo-EM micrographs were collected at National Cryo-EM Centre SOLARIS (Kraków, Poland) with the use of Titan Krios G3i microscope (Thermo Fisher Scientific, USA) operated at the accelerating voltage of 300 kV, magnification of 105k and corresponding pixel size of 0.86 Å/px. K3 direct electron detector was used for data collection in BioQuantum Imaging Filter (Gatan) setup with 20 eV slit enabled. The K3 detector was operated in counting mode with physical pixel resolution. Imaged areas were exposed to 41.20 e-/Å2 total dose each (corresponding to ~16.45 e-/px/s dose rate measured on vacuum). The images were acquired with EPU 2.10 software at under-focus optical conditions with a defocus of −3 µm. Scaling and cropping were done in Image J.

### 3.7 Nanoparticle tracking analysis

Concentration and size of secreted pTr - EVs were analyzed using NanoSight NS300 analyzer (Malvern Pananalytical). Before the measurement, samples were diluted in 0.2 µl-filtered DPBS without Ca^2+^ and Mg^2+^ (Lonza, USA) to reach particle concentration optimal for the measurement range of the instrument. Three 60 sec tracking repetitions of each sample were collected in syringe pump flow mode, using camera level of 13. Mean particle concentration and size distribution were calculated using Nanoparticle tracking analysis (NTA) Software ver. 3.4 (Malvern Pananalytical), with the threshold parameter set on 2.

### 3.8 Flow cytometry

pTr - EVs were stained with RNASelect dye (Thermo Fisher Scientific) at the final concentration of 5 µM and one of the following anti-human mouse monoclonal antibodies (5 µl per sample) conjugated with allophycocyanin (APC): anti-CD9 (clone MEM-61, Thermo Fisher Scientific), anti-CD63 (clone MEM-259; Thermo Fisher Scientific), anti-CD81 (clone 5A6; BioLegend) or the appropriate isotype control (mouse IgG1 k, Miltenyi Biotec). Prior to staining, RNASelect dye and antibodies were suspended in 0.2 µm-filtered DPBS (Lonza, USA) and centrifuged at 21 000 × g for 20 min at 4°C to eliminate potential debris and aggregates. Next, supernatants as staining buffers were mixed with EV samples in new tubes and incubated 20 min at 4°C. The analysis was performed with an Apogee A60-Micro-PLUS (Apogee Flow Systems). The percentage of gated positive events was calculated by Histogram software (Apogee Flow Systems). Control samples containing only staining buffers (without EVs) were also acquired to confirm the absence of potential background particles.

### 3.9 mRNA and miRNA real-time PCR

Total RNA was isolated using RNeasy Mini Kit (Qiagen, Germany) according to the standard manufacturer protocol and stored at −80°C for further real-time PCR (qRT-PCR) analysis. Gene expression was assessed using TaqMan Gene Expression Assays (Supplementary Table 3), TaqMan RNA–to Ct1-Step Kit (Thermo Fisher Scientific, USA) and Applied Biosystems Fast Real-time PCR system 7900HT according to the manufacturer’s protocol (Thermo Fisher Scientific, USA). In each qRT-PCR reaction (performed in duplicates), 15 ng of total RNA was used as a template. Negative controls without a template were performed in each run. The expression values were calculated and normalized to reference genes showing the best stability, calculated by NormFinder [29]. The geometric mean of two reference genes (*ACTB1* and *GAPDH*) with stability 0.106 for conceptuses/trophoblasts. For pTr cells, *GAPDH* was used (stability 0.028).

Small RNAs were isolated from pTr cells after miRNA delivery using a miRVana miRNA isolation kit. To confirm transfection efficiency, two-step real-time PCR was performed according to manufacturer instruction using TaqMan MicroRNA Assays (Life Technologies, USA), and 10 ng of total RNA was used for the reverse transcription (RT) using MultiScribe Reverse Transcriptase and RT primers specific for each miRNA. For qPCR reaction, 2.6 μL of cDNA was used along with TaqMan Universal Master Mix II and miRNA probes. Amplification was performed with denaturation for 10 min at 95 °C, followed by 45 cycles of 15 sec at 95 °C and 60 sec at 60 °C. Negative controls were performed in each run, without RNA or reverse transcriptase added to the reaction.

### 3.10 *In silico* screening of miRNA-mRNA binding motifs

To identify miRNA-mRNA binding sites, the sequences ranging from 5000 bp upstream and 200 bp downstream of the transcription starting site (TSS) for tested genes were extracted using biomaRt R package [30]. The regions were scanned for miR-125b-5p (CCCUGAG - seed sequence) matching 5 site types, i.e.: offset 6mer, 6mer, 7mer-A1, 7mer-m8 and 8mer using custom Python script. To compute the minimal free energy (mfe) of miRNA-target duplexes, RNAhybrid was used [31].

### 3.11 Statistical analysis

Statistical analysis was carried out in GraphPad Prism 9.4.1 (GraphPad Software Inc., USA). Effects were considered significant at p1<10.05. Logarithmic transformation of the data was performed for samples without a normal distribution (Shapiro-Wilk test). Data are presented as mean ± standard error of the mean (SEM), unless specified otherwise, as indicated in each figure legend. Statistical tests were selected based on the experimental set up and included: one-way ANOVA followed by Tukey’s post hoc test, unpaired T-test and paired T-test. The Pearson correlation coefficient (PCC) was used for quantitative analysis of immunolocalization. Sample sizes and other statistical details are indicated in the figures/figure legends and text.

## 4. RESULTS

### 4.1 Conceptus development and implantation are accompanied by changes in abundance of EV biogenesis and trafficking machinery

Our recent studies have shown that EV biogenesis and trafficking change at the embryo–maternal interface upon the arrival of the embryo into the uterus and that endometrial genes of the EV biogenesis pathway respond to embryonic signals [26]. For these reasons we decided to test whether developing porcine conceptuses changed their EV biogenesis and trafficking pathways to cope with the demands of robust development and implantation. To this end, three important time points during conceptus development and implantation were chosen [32] — maternal recognition of pregnancy and embryo apposition (days 11–12, from tubular to filamentous conceptuses), initiation of embryo attachment (days 15–16, elongated conceptuses), and full embryo attachment (days 17-19, elongated). First, *VPS28* and *VPS37B*, which belong to the ESCRT-I subunit [33], were examined. The highest abundance of *VPS28* was observed on days 15–16, with a significant drop afterward, while *VPS37B* levels gradually dropped between the tested days of pregnancy (Fig. 1A). Next, members of the ESCRT-II subunit responsible for the sorting of ubiquitinated cargo into MVBs were tested (i.e., *VPS22*, *VPS25* and *VPS36*) [34]. All tested ESCRT-II subunit genes showed the same pattern of expression with the higher mean levels on days 15–16 and significant decrease on days 17–19 (Fig. 1B). The third group of analyzed genes consisted of the coding ESCRT complex– associated proteins –*Alix*, *VPS4A,* and *VPS4B* [35], as well as vesicle–associated membrane protein 8 (*VAMP8*), which is in charge of EV secretion and release [36]. *Alix*, responsible for the abscission stage of cytokinesis during ILV formation, and *VPS4B,* which recognizes membrane-associated ESCRT-III assemblies and catalyzes their ATP-dependent disassembly [37] showed almost constant abundance during pregnancy. On the other hand, *VPS4A*, which redistributes the ESCRT-III components back to the cytoplasm [38], and *VAMP8* showed the highest mean levels on days 11–12, however, they reached the lowest levels in the subsequent days of pregnancy (i.e., days 15–16 for *VPS4A* and days 17–19 for *VAMP8*; Fig. 1C). The last group of genes analyzed code small RAB GTPases (*Rab11A, Rab11B, Rab7B, Rab8B,* and *Rab27A*), which are mainly involved in the intercellular trafficking of vesicles, MVB maturation, and vesicle secretion [39]. Only *Rab11B* showed changing abundance, gradually increasing between days 11–19 (Fig. 1D). These results suggest that the expression patterns of EV biogenesis and trafficking machinery vary in developing conceptuses dependent on the stage of implantation.

**Figure 1.**
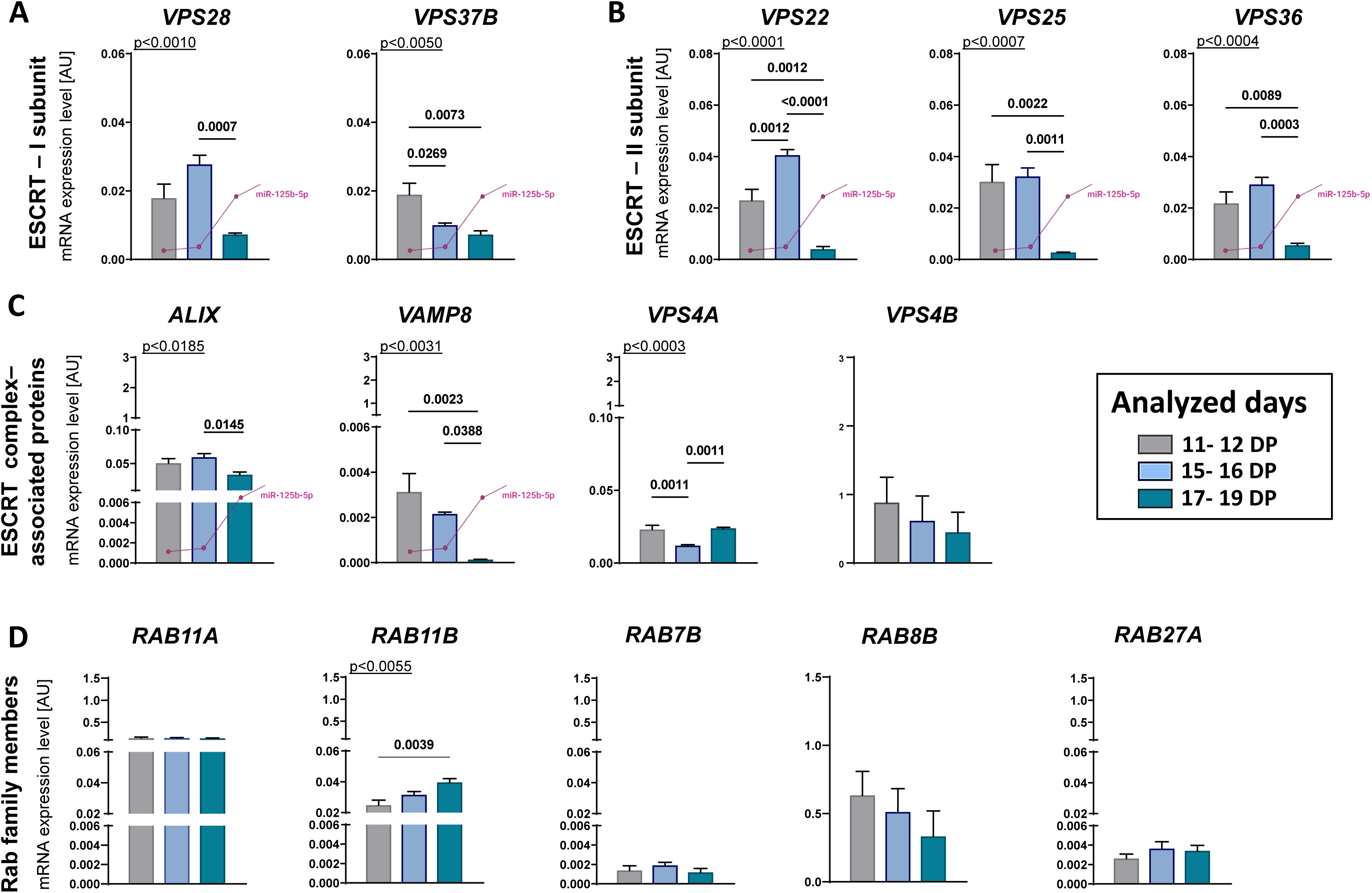
EV biogenesis and secretion pathways in conceptuses during early pregnancy. mRNA expression of **(A)** ESCRT-I complex proteins: VPS28, and VPS37b; **(B)** ESCRT-II complex proteins: VPS22, VPS25, and VPS36; **(C)** ESCRT complex–associated proteins: Alix, VAMP8, VPS4A, and VPS4B; and **(D)** Ras-related proteins: RAB11A, RAB11B, RAB7, RAB8B, and RAB27A. Gene expression was normalized to geometric mean of *GAPDH* and *ACTB1* (AU), identified as the best reference by NormFinder algorithm. Data were analyzed using one-way ANOVA with Tukey’s multiple comparison post-hoc test and are shown as means + SEM (n = 6-8). Significant differences between days (11-12, 15-16, and 17-19) are indicated in the graph (p < 0.05). AU – arbitrary units, DP – days of pregnancy. Previously reported profile of expression for miR-125b-5p in porcine conceptuses/trophoblasts [18] is marked in purple only for genes showing inverse pattern of expression.

### 4.2 miR-125b-5p affects the expression of genes involved in EV biogenesis and trafficking

miR-125b-5p is present in conceptuses and can be transported in EVs present at the embryo– maternal interface [18,19]. Its existence in the uterine environment during early pregnancy led to a series of experiments exposing miR-125b-5p as a potent fast-acting regulatory mediator of EV biogenesis and release during endometrial response to embryonic signals [26]. Furthermore, the delivery of miR-125b-5p to pTr cells affected gene expression, leading to changes in trophoblast migration properties [20]. To gain a full view of miRNA-mediated events at the embryo–maternal interface, we decided to examine the involvement of miR-125b-5p in the regulation of EV biogenesis and trafficking pathways in trophoblast. First, we compared the expression profile of genes coding different steps of the ESCRT-dependent EV biogenesis pathway, its associated proteins, and Rabs with the known profile of miR-125b-5p, which increases in abundance after day 16 of pregnancy [18]. We anticipated that genes previously characterized by a drop in expression on days 17–19 of pregnancy could be alerted by miR-125b-5p. Using in silico analysis, we searched for nucleotide positions 2–8 of miRNA, known as the seed sequence, which play crucial role in binding miRNA to the target region [40]. We focused on the localization of the perfect match of 6 to 7 seed nucleotides with the corresponding nucleotides of the target sequence, causing functional interactions between miRNA and the target. These sites are known as canonical site types [41,42]. miRNA response elements for miR-125b-5p were found within porcine genes coding *VPS36* and *VPS37B* (Fig. 2A), as well as *VPS22, VPS25, VPS28, Alix,* and *VAMP8* (Supplementary Fig. 2A). Next, after efficient in vitro delivered of miR-125b-5p to pTr cells (Fig. 2E, F; Supplementary Fig. 2B), we examined the abundance of the target genes. We observed decreased levels of one ESCRT-I (VPS37; Fig. 2B) and all ESCRT-II complex members (VPS22, VPS25, VPS36; Fig. 2C), as well as the associated protein – VAMP8 (Fig. 2D).

**Figure 2.**
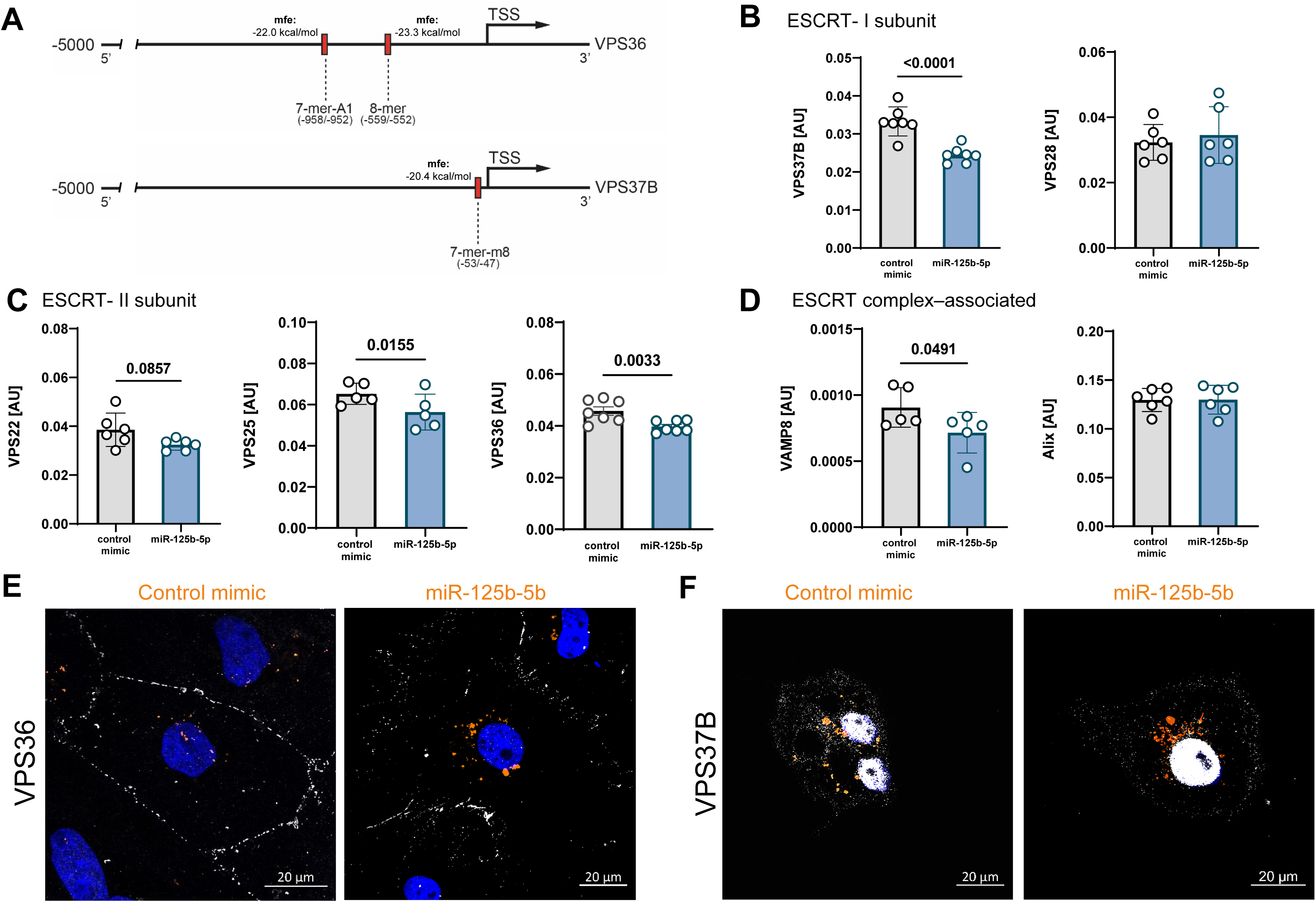
miRNA – mRNA target interactions. **(A)** Diagram showing positions of binding motifs for VPS36 and VPS37 targeted by canonical seeds presented along with minimum free energy (mfe) of miRNA-mRNA interaction. Expression of genes belonging to **(B)** ESCRT-I (VPS37B and VPS28), **(C)** ESCRT-II (VPS22, VPS25 and VPS36) complexes, and **(D)** ESCRT complex–associated proteins (VAMP8 and Alix). Gene expression was normalized to GAPDH (AU), identified as the best reference by distribution of VPS36 **(E)** or VPS37 **(F)** protein (grey scale) in pTr cells after delivery of control mimic (orange) or after miR-125b-5p (orange). mfe – minimal free energy; TSS – transcription start site.

Finally, we compared the cellular localization of internalized miR-125b-5p and its targets. We focused on VPS37B and VPS36 proteins (members of ESCRT-I and ESCRT-II, respectively), most responsive to miR-125b-5p delivery in vitro (Fig. 2) and showing opposite patterns of expression to miR-125b-5b in conceptuses between days 11–19 of pregnancy (Fig. 1). miR-125b-5p and negative control mimic (orange) were scattered throughout the cytoplasm, suggesting endosomal distribution. Initial observations showed the polarization of VPS36 at the pTr cell–cell borders in the control mimic and more scattered and diffused signal distribution after miR-125b-5p delivery (Fig. 2E). In contrast, VPS37B did not show treatment-specific patterns of signal distribution in the pTr cell cytoplasm (Fig. 2F). These results suggest that miR-125b-5p can act as a regulatory miRNA governing ESCRT-dependent EV biogenesis.

### 4.3 miR-125b-5p is efficiently delivered to pTr cells and internalized into endosomal trafficking

To further investigate the intracellular route of miRNA internalization after delivery to the pTr cells, we used the BacMam technology, based on the double-stranded DNA insect virus (baculovirus) as vehicle to efficiently deliver and express genes in mammalian cells. Late endosomes and MVBs throughout the endosomal trafficking pathways were reported to be involved in miRNA transition into recipient cells [43], specifically enhancing the miRNA-guided silencing by promoting recycling of RISC to engage with small RNA effectors and target RNAs more effectively [44]. MVBs are suggested as places where miRNA biogenesis and RISC-assembly pathways intersect [45]. To observe the subcellular localization of miRNA delivered to pTr cells, the BacMam-mediated fusion construct of Rab7 (marker of late endosomes/MVBs [46]) with emerald green fluorescent protein (EM-GFP) was used as a specific targeting tool for labeling late endosomes/MVBs (Fig. 3A). Rab7B antibody staining validated the proper labeling of the baculovirus construct (Fig. 3B, C, D). Super-resolution images and 2.5D models confirmed that after delivery miRNA was encapsulated in late endosomes/MVBs-like structures (Supplementary Material Movie 1; control, empty structures – Supplementary Material Movie 2). The results suggested that internalized miRNAs are transported via endosomal routes after delivery to pTr cells, pointing at the importance of endosomal and vesicle trafficking in miRNA intracellular processing at the embryo–maternal interface.

**Figure 3.**
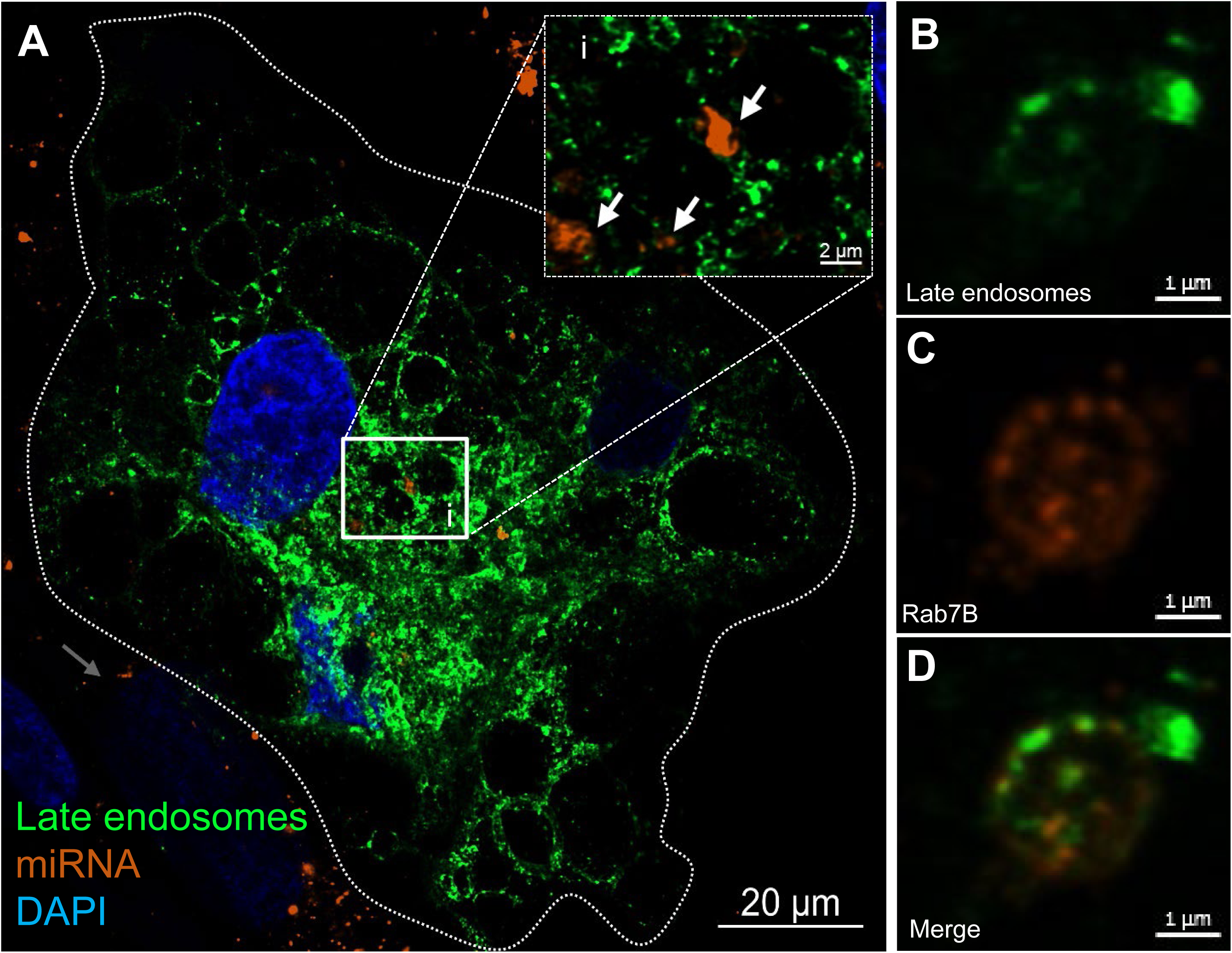
Endosomal route of miRNA internalization to late endosomes/MVBs. **(A)** Primary porcine trophoblast cells after miRNA delivery expressing fluorescently tagged - GFP RAB-7B GTPases. High resolution microscopy confirmed localization of internalized miRNA into late endosomes (higher magnification, white arrows). DAPI was used to visualize nucleus. **(C-E)** Specify of baculovirus construct revealed by RAB7B antibody staining.

### 4.4 CD63 and VPS36 showed spatial-dependent colocalization and correlation after miRNA delivery

Next, we verified the involvement of miR-125b-5p in the modulation of EV trafficking, focusing on the localization of EV-related proteins in pTr cells. VPS36 (miR-125b-5p target and marker of ESCRT-II complex present in endosomes/MVBs [47]) and CD63 (marker of EVs [48]) were selected based on our recent studies pointing to embryonic signals as triggers of EV trafficking in endometrial luminal epithelial cells [26]. Regions of interest (ROIs) were set based on the overlapping localization of both proteins present in the highly visible and well-shaped cell borders (white) in comparison to the whole cell (Fig. 4A). Identified miR-125b-5p–VPS36 interplay suggested that the miRNA delivered to pTr cells impairs the abundance of VPS36, involved in EV biogenesis, as required for the maturation of late endosomes/MVBs [49]. We anticipated that miR-125b-5p would also indirectly affect CD63 distribution, as an indication of the accumulation in MVBs and EV secretion [50]. Thus, we measured changes in VPS36 and CD63 fluorescent signal intensities, their cellular localization (cell–cell borders vs. whole cell) and colocalization. Analysis of the pTr cell–cell borders revealed three possible types of signal distribution profiles – VPS36^+^, CD63^+^, and VPS36^+^/CD63^+^ (Fig. 4B, C). After miRNA delivery, decreased signal intensity of VPS36 at the cell–cell borders (Fig. 4D), but not for the whole cells (Fig. 4G) was noted. Strikingly, no matching pattern was observed for CD63 signal intensity, which was not affected by miRNA delivery at the cell–cell borders (Fig. 4E), but increased in the whole cells (Fig. 4H). The colocalization coefficient analysis also revealed differences between cell–cell borders and the whole cells after treatment. At the cell–cell borders, a weak positive association between CD63^+^ and VPS36^+^ was identified for the control mimic, suggesting the ongoing release of EVs from MVBs as signals from CD63^+^ and VPS36^+^ were barely colocalized. After the miR-125b-5p delivery association became moderate, miRNA delivery led to pronounced MVB trafficking at cell–cell border (Fig. 4F). For the whole cells, this effect was no longer observed. A moderate correlation was noticed for the control mimic, while miR-125b-5p delivery led to weak, negative correlation between VPS36^+^ and CD63^+^, meaning decreased MVB trafficking (Fig. 4I). These differences in the cellular distribution of CD63^+^ and VPS36^+^ signals after miRNA delivery showed a pattern that we were able to detect only within well-defined regions located on the preserved cell–cell borders. Together these results point to the role of miR-125b-5p in VPS36-mediated changes in vesicular trafficking in pTr cells.

**Figure 4.**
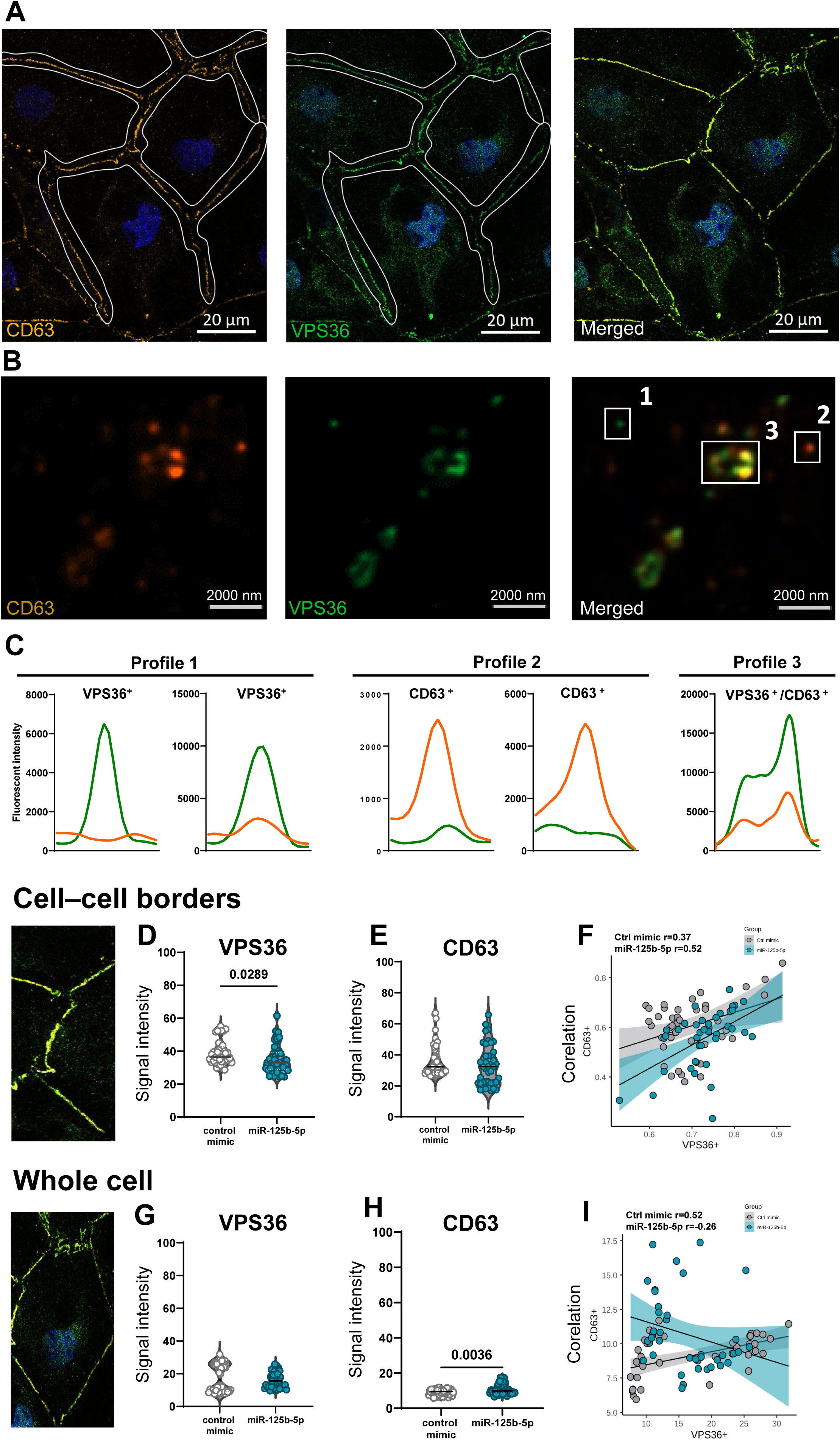
Spatial-dependent colocalization and correlation of VPS36 and CD63 after miRNA delivery to pTr cells. **(A)** Identification of cell-cell contact areas (regions of interest, ROIs) was performed based on the overlapping CD63 (CY3, orange) and VPS36 (Alexa 488, green) signals at cell-cell contact areas. Nucleus was stained with DAPI. Merged images show an example of pTr cell colony. **(B)** Profiles for CD36 and VPS36 colocalization at cell-cell border (ROIs). **(C)** Schematic representation of VPS36+, CD63+, and VPS36+/CD63+ profiles, based on their signal intensity. Signal intensities for VPS36 **(D)** and CD63 **(E)** at cell-cell borders after treatment with control mimic or miR-125b-5p. Data are presented as violin plots with individual samples and median indicated (unpaired T-test, p < 0.05). **(F)** Scatter plots of CD63+ and VPS36+ pixels at cell-cell borders corresponding to the colocalization events indicate weak correlation and linear relationship for control mimic (r = −0.37, p = 0.018) and moderate correlation and linear relationships for miR-125b-5p (r = 0.52, p = 0.001). Whole cell signal intensities for VPS36 **(G)** and CD63 **(H)** after mimic or miR-125b-5p delivery. Data are presented as violin plots with individual samples and median indicated (unpaired T-test, p < 0.05). **(I)** Scatter plots of CD63+ and VPS36+ pixels in whole cell analysis corresponding to the colocalization events indicate moderate correlation and linear relationship for control mimic (r = 0.52, p = 0.001) and negative correlation and non-linear relationships for miR-125b-5p (r = −0.26, p = 0.103). Experiments were performed in four solid biological replicates. Pearson correlation coefficient was performed (p < 0.05).

### 4.5. miR-125b-5p delivery to pTr cells promote CD63 EVs secretion

Given that miR-125b-5p affects the expression of genes involved in EV biogenesis and changes vesicular trafficking in pTr cells, we hypothesized that it can also influence the composition of EV subpopulations and their secretion. Following the stepwise centrifugation of media collected after treatment with the control mimic or miR-125b-5p, EV pellets were used for the characterization of secreted EV subpopulations (Fig. 5A). Cryo-EM revealed the presence of round-shaped EVs with a clear lipid bilayer (Fig. 5B). In general, no differences between the concentration (Fig. 5C) and size of EVs released by pTr cells after treatment with miR-125b-5p (mean: 131.4 ± 2.38 nm) and control mimic (mean: 128.3 ± 3.04 nm) were noted (Fig. 5D). However, analysis of the particle size distribution within the sample showed significant differences. The cumulative parameters D10, D50, and D90 indicating that 10%, 50% or 90% of the particles, respectively, had a diameter less than or equal to the given value; D10 and D50 were shown to be higher in EVs collected after treatment with miRNA-125b-5p (Fig. 5D). We expected that miRNA delivery to pTr cells would affect ESCRT-dependent pathways and consequently change the EV subpopulation ratio, as other biogenesis pathways can be activated. High-resolution flow cytometry was used to determine the CD9^+^, CD63^+^, and CD81^+^ subpopulations of the EVs. The characteristic profile of tetraspanins was discovered – CD81^+^ > CD63^+^ > CD9^+^ (Fig. 5E, F). Strikingly, CD63-bearing EVs were more abundantly secreted by pTr cells after miR-125b-5p delivery (Fig. 5G). Taken together, these findings show that miR-125b-5p can affect EV subpopulations, leading to the increased release of CD63^+^ EVs.

**Figure 5.**
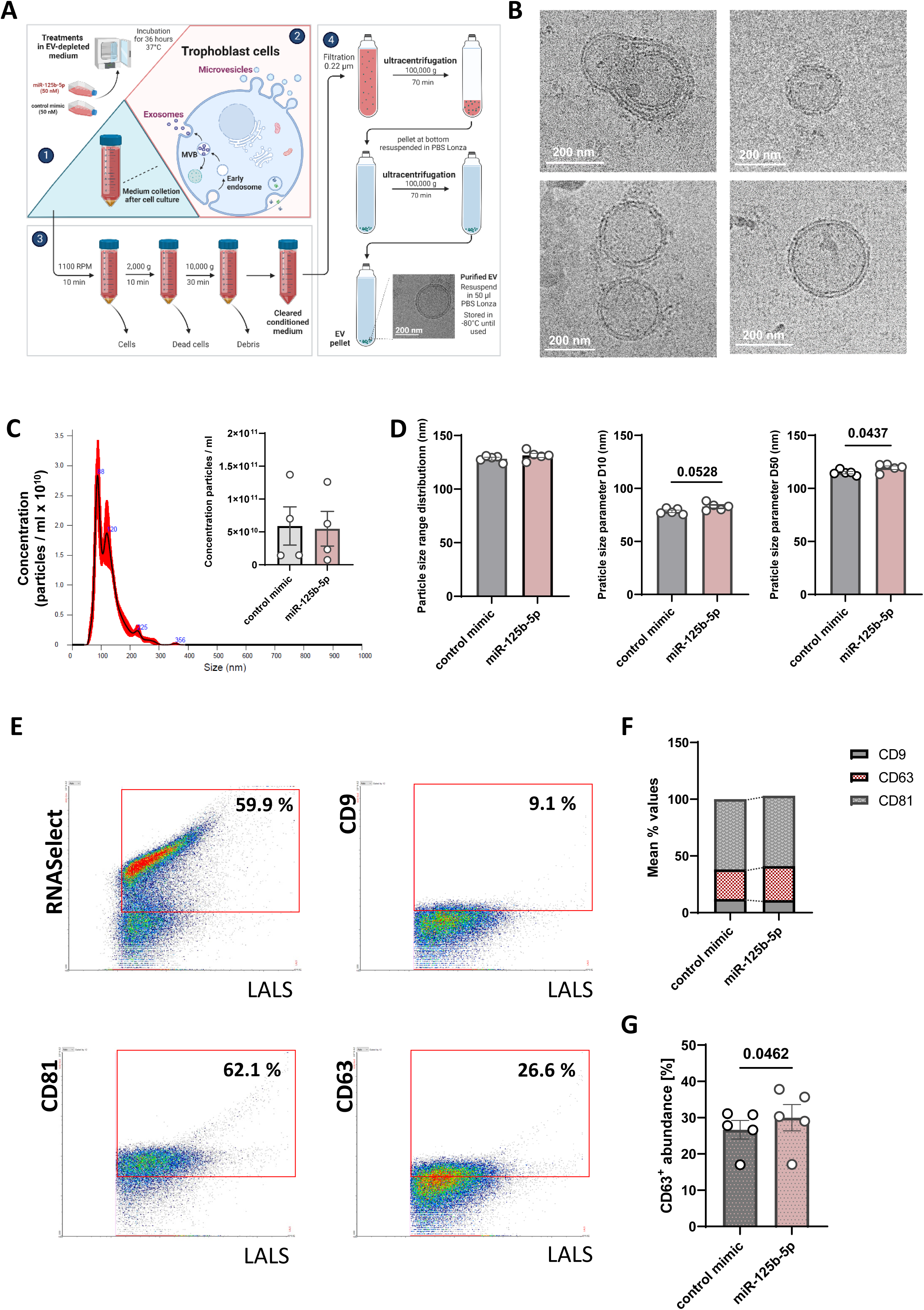
Subpopulations of EVs secreted by pTr cells. **(A)** Schematic diagram showing experimental design of stepwise centrifugation to obtain pTr-secreted EVs. **(B)** Cryo-EM images showing representative extracellular vesicles secreted by pTr cells. **(C)** Representative profile of particle distribution showing size and concentration of EVs derived from the pTr cells. **(D)** Particle size range distribution and particle size parameters (D10 and D50; paired T-test). **(E)** Representative plots of pTr-derived EVs stained with RNASelect dye and fluorescent antibodies (anti-CD63, -CD9, and -CD81). The percentage of objects positive for the analyzed EV marker is shown in red gates. LALS - large angle light scatter parameter, corresponding to the relative size of analyzed particles. **(F)** Surface presence of selected EV markers (CD63, CD9, CD81) between samples collected after either control mimic or miR-125b-5p. **(G)** Abundance of CD63 positive EVs secreted by pTr cells after delivery of control mimic or miR-125b-5p (paired T-test)

## 5. DISSCUSSION

The involvement of EVs in early pregnancy events has been highlighted for several mammalian species, including humans and domestic animals, such as the pig [51,52,5,19]. In recent years, EVs as transporters of miRNAs have also been considered as crucial for gene expression regulation at the embryo–maternal interface [17,18,20]. In our recent report, we showed that signals present in the uterine environment (e.g., non-coding RNAs and hormones) impact the EV biogenesis and trafficking at the maternal site of the dialog [26]. In this study, we focused on the conceptus and trophoblast to gain a full view of the EV-mediated embryo–maternal communication. We revealed the important role of miR-125b-5p in EV biogenesis and trafficking pathways. The inhibitory effect of miR-125b-5p on i) genes belonging to ESCRT-dependent biogenesis pathway and ii) the spatial distribution of MVBs containing the main EV marker – CD63 – was demonstrated. miR-125b-5p occurred as a negative regulator of VPS36 recruitment at pTr cell borders, accompanied by the increased release of CD63-bearing EVs. Here, we provide unique evidence for the miRNA-mediated mechanism governing EV biogenesis and release at the embryonic site.

ESCRT-mediated processes function in a wide variety of cellular and developmental processes. Four ESCRT multi-subunit protein complexes act cooperatively at specialized endosomes to facilitate the movement of specific cargoes [53,54]. We identified inverse expression patterns of miR-125b-5p and proteins belonging to ESCRT machinery – namely *VPS22, VPS25, VPS28, VPS36, VPS37B,* (ESCRT-I and ESCRT-II components), and Alix (ESCRT complex–associated proteins) – showing the highest level around days 15–16 of pregnancy and a significant drop on days 17–19. The observed results are consistent with our recent studies showing overlapping profiles for the same genes during pregnancy in porcine endometrium [26], suggesting common regulatory pathways involving embryonic signals and non-coding RNAs. Among ESCRT complex–associated proteins, VAMP8, involved in MVB exocytosis [55], showed the highest abundance on days 11–12, when maternal recognition of pregnancy in pigs occurs. This can be a sign of the increasing secretory activity of conceptuses, leading to the release of embryonic signals for the maternal recognition of pregnancy [56]. On the other hand, an abundance of RAB11B, which regulates the slow recycling of membrane components from endosomal compartments [57] and is engaged in the secretion of EVs enriched in, e.g., flotillin [], was gradually increasing throughout the following days of pregnancy, when the attachment of the trophectoderm to the luminal epithelium is becoming stronger. Firm attachment of the trophectoderm involves integrins, a large family of proteins involved in the attachment, migration, invasion, and control of cellular function [58,59]. Various molecular pathways are polarization, including EV biogenesis and the trafficking of molecules [26]. Endosomal trafficking regulates the polarization of various integrins and their receptors suggesting involvement of Rabs in cell polarization [60]. There is a chance that the Rab-mediated recycling of integrins and their ligands might be important for trophoblast cell motility, as shown for breast cancer cells [61].

Intercellular transport is crucial for maintaining homeostasis within the cell, which includes developing conceptuses responding to physiological signals exchanged at the embryo–maternal interface. Vesicular transport as a major cellular activity plays a central role in the trafficing of molecules between different membrane-enclosed compartments of the secretory pathway [62]. According to previous reports implying that internalized miRNAs can be found on different levels of endosomal trafficking pathways, including MVBs and endosomes [63], we confirmed the delivery of miR-125b-5p into endosomal pathways in primary pTr cells. This also supports findings showing that small RNAs can be secreted from cells through the EV-mediated pathway [64,18]. Carried by EVs, miR-125b-5p is considered to be an important modulator of porcine trophoblast cell function, especially given its involvement in epitheliochorial placentation in pigs [20]. Recently, we showed that miR-125b-5p impacts ESCRT-dependent EV biogenesis pathway in primary porcine luminal epithelial cells – the first layer that receives signals from developing conceptuses. Furthermore, the miR-125b-5p–dependent down-regulation of ESCRT pathway machinery in luminal epithelial cells was abolished by embryonic signaling [26].

ESCRT-dependent pathway relies on interaction of all complexes. ESCRT-0 and ESCRT-I, involved in cargo sorting, are not sufficient for MVB generation, as the ESCRT-II–mediated initiation of ESCRT-III assembly is mandatory to complete this pathway [65]. The overexpression of miR-125b-5p in primary pTr cells led to the down-regulation of target genes belonging to ESCRT-dependent EV biogenesis pathway, especially the ESCRT-II complex. Previously conducted studies have shown that ESCRT-II, based on interactions between RILP (Rab-interacting lysosomal protein) and VPS22 and VPS36, change protein sorting into the MVBs and their morphology [66]. Thus, miRNA-mediated down-regulation of the ESCRT-II complex is likely to affect MVB trafficking and later, the EV secretion profile. Indeed, miR-125b-5p delivery to pTr cells changed the distribution of VPS36 from polarized at the cell borders to more diffuse and lacking polarization, which was accompanied by an altered tetraspanin profile in the secreted EVs. Several studies have independently found that ESCRT machinery regulates cell polarization and migration [67]. The adverse impacts of miR-125b-5p on ESCRT-dependent machinery and VPS36 recruitment to MVBs during cell polarization and signal transduction have physiological relevance in porcine pregnancy. At the time when miR-125b-5p expression increases in trophoblasts [18], cell migration is decreased, and the attachment phase has already been initiated [32].

Building a mechanistic understanding of EV-based embryo–maternal crosstalk requires studies focusing on both sides of this dialog. Our observations of VPS36 protein (as an ESCRT-II member and MVB marker) and EV-positive tetraspanin – CD63 distribution in trophoblast cells suggested that miRNA-induced targeting may be closely related to the generation and distribution of MVBs and other endosomes designated for MVB trafficking. Consistent with our previous studies on the maternal site of the dialog – the endometrium [26] – overlapping colocalization of VPS36 and CD63 at the cell–cell borders was also indicated for pTr cells. Embryonic signals (mainly E2), acting as modulators of EV secretion, led to an increase in VPS36 intensity at the endometrial luminal epithelium cell–cell borders and resulted in a loss of correlation between CD63^+^ and VPS36^+^ puncta. In contrast, at the trophoblast cell borders, the VPS36 protein level decreased after miR-125b-5p delivery. This was accompanied by a higher incidence of CD63^+^ and VPS36^+^ puncta colocalization, a sign of disturbed MVB trafficking and EV secretion. This pattern was spatial-dependent, seen only at the cell–cell borders, highly active location of cell-to-cell communication during pregnancy. Whole cell analysis revealed increased signal intensity for CD63. This is a common observation for most cell types, as the majority of CD63 is present at endosome-associated membranes rather than surface plasma membranes [68]. Within MVBs, CD63 is approximately seven times enriched than within ILVs [68]. Nevertheless, increased surface localization of CD63 is required in multiple signaling pathways governing cell proliferation, motility and adhesion as well as protein trafficking [68,69].

EV-mediated intercellular communication is based on the delivery of cargo to recipient cells; however, cells have the ability to constantly release various types of EV subpopulations with different morphology and cargo [70]. Considering the complex nature of miRNA regulatory mechanisms, we suspected that the identified miR-125b-5p–ESCRT interplay may cause changes in EV generation and the distribution at embryo–maternal interface. In fact, the repression of VPS36 recruitment to MVBs and the increased CD63 intensity in the whole cell could be an indication of VPS36 inhibition by miR-125b-5p. ESCRT-dependent and ESCRT-independent pathways coexist in numerous biological processes and are responsible for ensuring constant EV biogenesis and release [9,71,72,73]. Difficulty in finding the perfect blocker of the formation and release of vesicles hints at the possible overlapping of pathways to backup EV-mediated communication [74,75]. Therefore, the activation of ESCRT-independent pathways after miRNA delivery is possible, especially given that ceramide [68].

Recognizing the basic mechanisms underlying EV heterogeneity is essential to determining functional role of EVs. Analysis of the percentage population ratio revealed that miR-125b-5p delivery to pTr cells leads to the slightly increased size of released EVs. This finding is supported by our gene expression and colocalization studies showing the miRNA-mediated inhibition of ESCRT-dependent endosomal/MVB trafficking pathway, which is mainly responsible for the generation of small EVs [76]. Importantly, we also found that in response to miR-125b-5p, pTr cells begin secreting more CD63^+^ EVs. The main CD63 partners include integrins, other tertraspanins and surface receptors [8,68]. The role of integrins in the processes of trophoblast attachment as well as invasive and noninvasive implantation has been widely recognized in many species, including pigs [77,78]. It is likely that the miR-125b-5p–mediated inhibition of ESCRT pathway after day 16 of pregnancy may lead to the enrichment of CD63 in EVs released by trophoblasts to the uterine cavity, consequently providing opportunities for the proper establishment of pregnancy.

## 6. CONCLUSIONS

We favor the hypothesis that miRNA presence in the uterine environment is a key element of ongoing EV-mediated communication between the mother and developing conceptus, leading to the generation and secretion of specific subpopulations of EVs. This study is the first to show the regulatory effects of miR-125b-5p on ESCRT-mediated biogenesis and the further secretion of EVs by trophoblast cells at the time when the crucial steps of implantation are taking place. This highlights the power of miRNAs to govern processes assuring pregnancy success. Further studies should explore ESCRT-independent EV biogenesis pathways to gain a full view on EV-mediated communication at the embryo–maternal interface.

## ACKNOWLEDGEMENTS

The authors are grateful to M. Romaniewicz, M. Sikora, K. Gromadzka-Hliwa, M. Blitek, and J. Kłos from the Institute of Animal Reproduction and Food Research, Polish Academy of Sciences (IARFR, PAS) for either excellent technical assistance laboratory or help in care and handling of animals. We are also grateful to Dr K. Myszczynski (IARFR, PAS) and Dr D. Panas (IARFR, PAS) for support in statistical and bioinformatics analysis, E. Wator (MCB, Jagiellonian University) for help in cryo-EM imaging, and Y. Heifetz (The Hebrew University of Jerusalem) for critical comments during data interpretation and manuscript preparation.

## AUTHOR’S CONTRIBUTION STATEMENT

MMG designed and performed experiment, collected, analyzed, and interpreted data; prepared graphics, drafted the manuscript and participated in the preparation of its final version. MMG and KJW performed the confocal microscopy imaging and analyzed the data. EK was responsible for flow cytometry and nanoparticle tracking analysis. MR performed cryo-EM imaging. EZS provided access to equipment (Apogee A60-Micro-PLUS and NanoSight NS300) and methodology for nanoparticle analysis. MMK conceived and supervised the study, designed experiments, analyzed, and interpreted data and was responsible for the final version of the manuscript. Graphics were created with BioRender.com.

## DATA AVAILABILITY STATEMENT

All data generated or analyzed during this study are included in this published article or in the supplementary materials.

## ETHICAL STATEMENT

All procedures involving animals were conducted in accordance with the national guidelines for agricultural animal care in compliance with EU Directive 2010/63/UE.

## SUPPLEMENTARY MATERIALS

**Supplementary Table 1.** Assays used in miRNA in vitro delivery.

**Supplementary Table 2.** Primary and secondary antibodies used in immunofluorescent staining.

**Supplementary Table 3.** TaqMan gene expression assays used in real-time PCR.

**Supplementary Figure 1.** Control staining of pTr cells in culture performed without primary antibodies. Merged pictures with: **(A)** Red channel (Cy3) and **(B)** green channel (Alexa Fluor 488) compared with nucleus stained with DAPI and actin filaments stained with F-actin antibodies (Alexa Fluor 488 or 594).

**Supplementary Figure 2. (A)** Diagram showing positions of binding motifs for VPS28, VPS22, VPS25, VAMP8, and ALIX targeted by canonical seeds presented along with minimum free energy (mfe) of miRNA-mRNA interaction. **(B)** Delivery of miR-125b-5p to pTr cells measured using real-time PCR. Data were analyzed using paired t-test (n=3 animals).

**Supplementary Movie 1.** Late endosomes with internalized miRNA

**Supplementary Movie 2.** Late endosomes without internalized miRNA

