## Supplementary Figure 1 for "miR-125b-5p impacts extracellular vesicle biogenesis, trafficking, and EV subpopulation release in the porcine trophoblast by regulating ESCRT-dependent pathway"

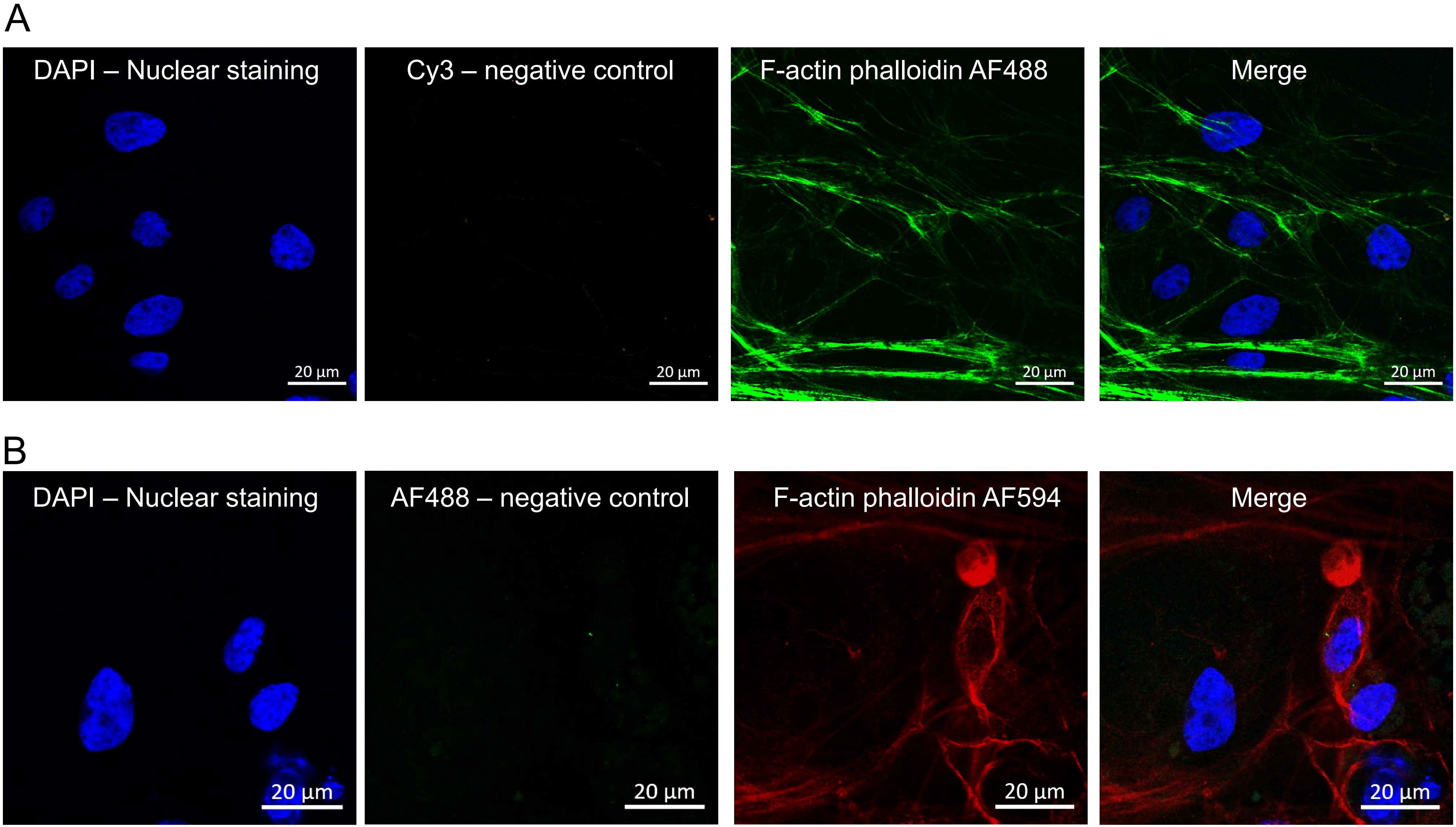


**Supplementary Figure 1.** Control staining of pTr cells in culture performed without primary antibodies. Merged pictures with: (A) Red channel (Cy3) and (B) green channel (Alexa Fluor 488) compared with nucleus stained with DAPI and actin filaments stained with F-actin antibodies (Alexa Fluor 488 or 594).
