## Supplementary Figure 2 for "miR-125b-5p impacts extracellular vesicle biogenesis, trafficking, and EV subpopulation release in the porcine trophoblast by regulating ESCRT-dependent pathway"

A

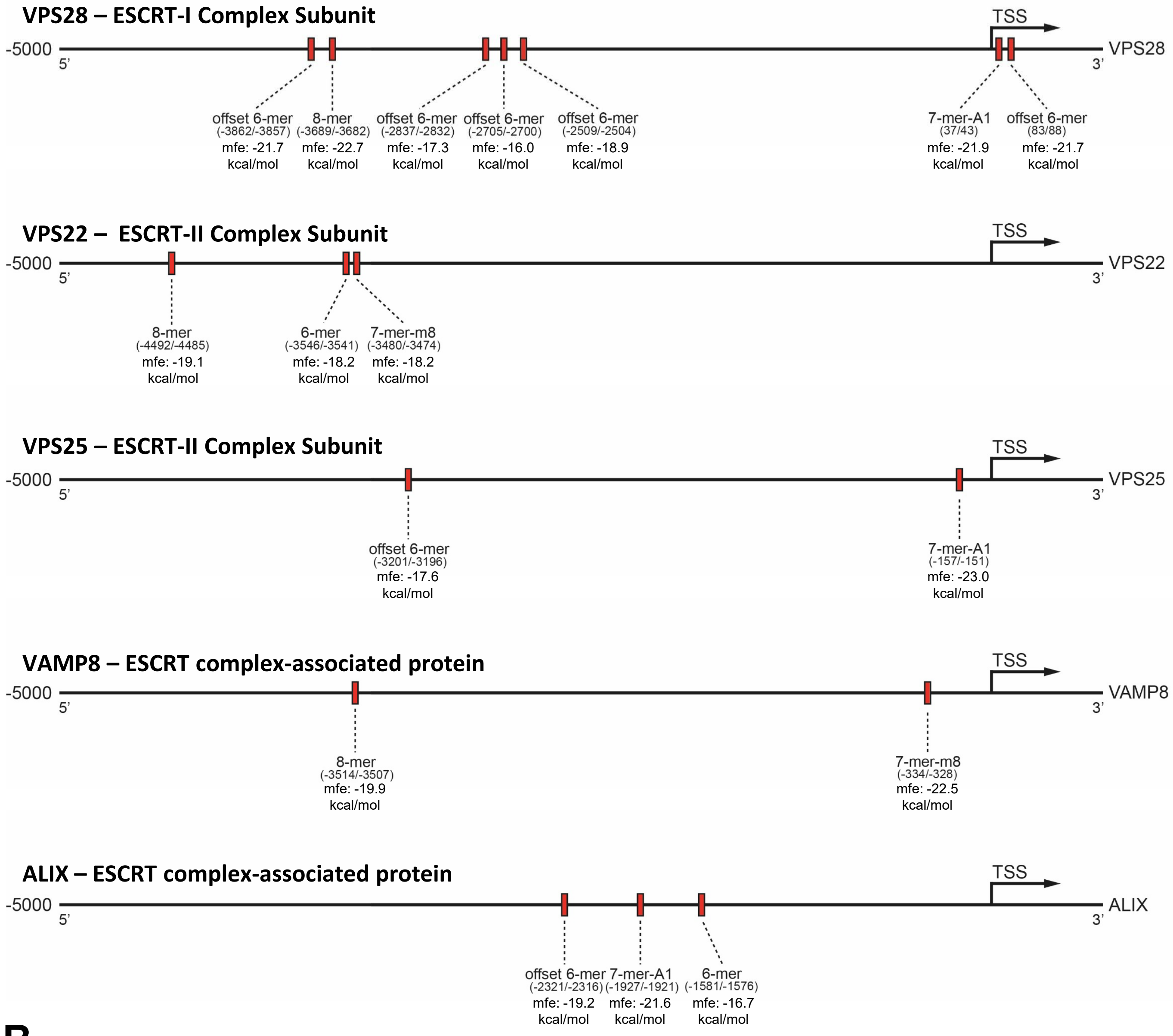


B

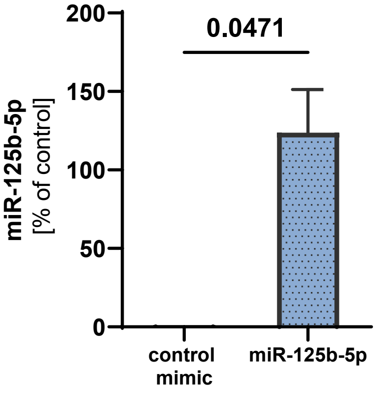


Supplementary Figure 2. (A) Diagram showing positions of binding motifs for VPS28, VPS22, VPS25, VAMP8, and ALIX targeted by canonical seeds presented along with minimum free energy (mfe) of miRNA-mRNA interaction. (B) Delivery of miR-125b-5p to pTr cells measured using real-time PCR. Data were analyzed using paired t-test (n=3 animals).
